# Limited genetic diversity of *bla*_CMY-2_-containing IncI1-pST12 plasmids from Enterobacteriaceae of human and broiler chicken origin in the Netherlands

**DOI:** 10.1101/2020.07.09.195461

**Authors:** Evert den Drijver, Joep J.J.M. Stohr, Jaco J. Verweij, Carlo Verhulst, Francisca C. Velkers, Arjan Stegeman, Marjolein F.Q. Kluytmans-van den Bergh, Jan A.J.W. Kluytmans, i-4-1-Health Study Group

## Abstract

Distinguishing epidemiologically related and unrelated plasmids is essential to confirm plasmid transmission. We compared IncI1-pST12 plasmids from both human and livestock origin and explored the degree of sequence similarity between plasmids from Enterobacteriaceae with different epidemiological links. Short-read sequence data of Enterobacteriaceae cultured from humans and broilers were screened for the presence of both a *bla*_CMY-2_ gene and an IncI1-pST12 replicon. Isolates were long-read sequenced on a MinION sequencer (OxfordNanopore Technologies). After plasmid reconstruction using hybrid assembly, pairwise single nucleotide polymorphisms (SNP) were determined. The plasmids were annotated, and a pan-genome was constructed to compare genes variably present between the different plasmids. Nine *Escherichia coli* sequences of broiler origin, four *Escherichia coli* sequences and one *Salmonella enterica* sequence of human origin were selected for the current analysis. A circular contig with the IncI1-pST12 replicon and *bla*_CMY-2_ gene was extracted from the assembly graph of all fourteen isolates. Analysis of the IncI1-pST12 plasmids revealed a low number of SNP differences (range of 0-9 SNPs). The range of SNP differences overlapped in isolates with different epidemiological links. One-hundred and twelve from a total of 113 genes of the pan-genome were present in all plasmid constructs. NGS-analysis of *bla*_CMY--*2*_-containing IncI1-pST12 plasmids isolated from Enterobacteriaceae with different epidemiological links show a high degree of sequence similarity in terms of SNP differences and the number of shared genes. Therefore, statements on the horizontal transfer of these plasmids based on genetic identity should be made with caution.

## Introduction

Antimicrobial resistance in Gram-negative bacteria is a worldwide growing public health problem [1, 2]. The gut is an important reservoir for resistant Gram-negatives, both in humans and livestock [3, 4]. Antimicrobial resistance in livestock has been suggested as potential source for resistance in humans, with a growing number of studies published on this potential transmission route for antimicrobial resistance mechanisms in Gram-negative bacteria [5–7]. AmpC beta-lactamase-production is an example of these mechanisms, causing 3^rd^ generation cephalosporin resistance in Gram-negative bacteria [8]. Plasmids are an important vector for antimicrobial resistance dissemination with genes for various resistance mechanism (e.g. AmpC beta-lactamase genes) being located on these mobile genetic elements. Incompatibility group I1 (IncI1) plasmids of the plasmid sequence type (pST) 12 have been associated with the spread of *bla*_CMY-2_, the most common AmpC beta-lactamase gene [9–11]. Recent studies show that the sequence of IncI1 plasmids is highly conserved [12–15]. Most studies to date are based on short-read sequence data [13, 14]. However, it remains challenging to study plasmid transmission using short-read sequencing data alone. Repeated sequences, often shared between plasmid and chromosomal DNA, hinder the assembly of the bacterial genome from short-read data, often resulting in contigs of which the origin, either plasmid or chromosomal, cannot be resolved [16]. This limits the interpretation of plasmid transmission by not providing accurate prediction of the total plasmid sequence. Recently, a combination of short- and long-read sequence data has provided the possibility for more accurate analysis, such as shown in a recent study on IncI1 plasmids of pST 3 and pST7 [15]. However, combined short- and long-read sequencing data of IncI1-pST12 plasmids from human and livestock origin is still absent. Transmission of antimicrobial resistant bacteria within and between domains is predominantly based on the comparison of bacterial chromosome. However, when only typing the bacterial chromosome, transmission of resistance gene containing plasmids can go undetected. Data on the within and between domain (human vs livestock) transmission of plasmids containing the AmpC beta-lactamase genes is limited. Accurately distinguishing related from non-related plasmids based on molecular characteristics (e.g. number of SNP differences) is essential for using sequence data to detect plasmid transmission. We hypothesize that a combination of short- and long-read sequence data of *bla*_CMY-2_ containing IncI1-pST12 plasmids reveal highly conserved plasmid sequencing, which complicates distinguishing plasmid transmission between epidemiologically related and unrelated isolates. The objective of the current study is to determine the relatedness between IncI1-pST12 plasmids of epidemiologically related and unrelated Enterobacteriaceae isolates from humans and livestock and we explore the possibility of accurately distinguishing related from unrelated samples based on plasmid sequencing data alone.

## Methods

### Collection of isolates

#### AmpC E. coli isolates from i-4-1-Health Dutch-Belgian Cross-border Project

As part of the i-4-1-Health project, human and broiler samples were collected as described by Kluytmans-van den Bergh et al [17]. After vortexing, the swab was plated on Blood Agar plate (growth control, performed since 2011) and the liquid Amies eluent was inoculated in selective tryptic soy broth (TSB) and incubated for 18–24 hours (35–37°C). Broths were subcultured on a AmpC selective MacConkey agar containing on one side cefotaxime 1 mg/L, cefoxitin 8 mg/L and on the other side ceftazidime 1 mg/L, cefoxitin 8 mg/L (Mediaproducts, Groningen) [18]. For all oxidase-negative isolates that grew on either side of the selective agar plates, species identification was performed by MALDI-TOF (bioMérieux, Marcy l’Etoile, France). Susceptibility testing was performed using Vitek 2 (bioMérieux, Marcy l’Etoile, France). The presence of AmpC in all oxidase-negative isolates was phenotypically confirmed using the D68C AmpC & ESBL Detection Set (Mastdiscs, Mastgroup Ltd, Bootle United Kingdom) and interpreted according to manufacturer’s instructions. All phenotypically confirmed isolates were sequenced using a Illumina MiSeq sequencer (Illumina, San Diego, CA, USA). DNA isolation and sequencing was performed as described by Coolen et al [19]. De novo assembly and error-correction was performed using SPAdes version 3.9.1 [20].

#### AmpC E. coli isolates from Amphia prevalence screening

pAmpC containing *E. coli* isolates were selected from a prevalence screening which had been performed in the Amphia hospital described by Den Drijver et al [21]. Rectal swabs taken from hospital patients were pre-enriched using selective TSB and subsequently cultured on MacConkey agar plate containing cefotaxime (1 mg/L) or MacConkey double agar plate containing cefotaxime (1 mg/L) with cefoxitin (8 mg/L) one side and ceftazidime (1 mg/L) with cefoxitin (8mg/L) other side (Mediaproducts, Groningen, The Netherlands) [18].

WGS was performed in UMCG using MiSeq (Illumina, San Diego, United States) and assembled with CLC Genomics Workbench 9.0, 9.0.1 or 9.5.2 (Qiagen, Hilden, Germany) as was previously described in more detail by Kluytmans-van den Bergh et al [22].

#### pAmpC-encoding clinical isolates from Elisabeth-Tweesteden hospital

Suspected pAmpC containing *E. coli* isolates from blood cultures were selected retrospectively from our laboratory database based upon the presence of phenotype (FOX MIC > 8 mg/L and/or CTX MIC ≥ 1mg/L and/or CAZ MIC MIC ≥ 1mg/L. One *Salmonella enterica* serotype Kentucky isolate from a fecal sample was selected from our laboratory database based upon the presence of AmpC suspected phenotype (FOX MIC > 8 mg/L and/or CTX MIC ≥ 1mg/L and/or CAZ MIC MIC ≥ 1mg/L). The isolates were recultured from deep frozen samples on blood agar and identified using the MALDI-TOF MS (BD Diagnostic Systems, Sparks, MD, USA). Susceptibility testing was performed using Phoenix Automated Microbiology System (BD Diagnostic Systems, Sparks, MD, USA). The isolates were sequenced using a Illumina MiSeq sequencer (Illumina, San Diego, CA, USA). DNA isolation and sequencing was performed as described by Coolen et al [19]. De novo assembly and error-correction was performed using SPAdes version 3.9.1 [20].

### Whole-genome bioinformatics analysis of short –read sequencing data

The presence of acquired resistance genes was identified by uploading assembled genomes to the ResFinder web-service of the Center for Genomic Epidemiology (version 3.1) [23]. The presence of plasmid replicons and the typing of specific IncI plasmid was performed using pMLST (version 2.0) [24]. The genomes were selected based on a 100% match to *bla*_CMY-2_ and IncI-pST12. MLST typing was performed using MLST web-service of the Center for Genomic Epidemiology (version 2.0), and *fim* typing was performed using FimTyper (version 1.0), Center for Genomic Epidemiology [25, 26]. Whole-genome MLST (wgMLST) (core and accessory genome) was performed for all isolates using Ridom SeqSphere+, version 4.1.9 (Ridom, Münster, Germany). Species-specific wgMLST typing schemes were used as described previously [22]. The pairwise genetic difference between isolates of the same species was calculated by dividing the total number of allele differences by the total number of shared alleles, pairwise ignoring missing values. Genetic relatedness was determined using the thresholds for wgMLST-based genetic distance as described previously [22].

### Long-read sequencing and hybrid assembly

No more than two isolates of the same flock or patient with the same wgMLST-based genetic distance were selected for further long-read sequencing

All isolates were long-read sequenced on a MinION sequencer using the FLO-MIN106D flow cell and the Rapid Barcoding Sequencing Kit SQK RBK004 according to the standard protocol provided by the manufacturer (Oxford Nanopore Technologies, Oxford, United Kingdom). A hybrid assembly of long-read and short-read sequence data was performed using Unicycler v.0.8.4 [27].

### Plasmid analysis

The genomes created using the hybrid assembly were uploaded to the online bioinformatics tools ResFinder v.2.1, VirulenceFinder v.1.2 and PlasmidFinder v.1.2. (Center for Genomic Epidemiology, Technical University of Denmark, Lingby, Denmark) [23, 24, 28]. Circular components created by the hybrid assembly that were smaller than 1000kb and that contained an IncI1-pST12 plasmid replicon and a *bla*_CMY-2_ gene were extracted from the assembly graph using BANDAGE v0.8.1. [29]. All extracted plasmid components were annotated using Prokka v1.13.3 [30]. Using snippy v4.4.5 9 (https://github.com/tseemann/snippy) the number of single nucleotide polymorphisms (SNP) was determined between the extracted plasmid components using a *bla*_CMY-2_ gene containing IncI1-pST12 plasmid extracted from the GenBank (accession number: MH472638.1) as reference [12]. A pan-genome was constructed, and a gene-presence or absence was determined for all extracted plasmid components using roary v3.12 [31]. All extracted plasmids consisting of a single circular contig were aligned using GView 1.7 [32] and progressiveMAUVE v2.4.0. to detect possible rearrangements [33].

### Classification of pairwise comparisons

Pairwise comparisons of assembled plasmids were classified according to the known epidemiological link between the isolates: (i) same sample; (ii) same ward/flock, but different sample; (iii) same location (hospital or farm), but different ward/flock and sample; (iv) same domain (human or broiler), but different location, ward/flock and sample; and (v) no known epidemiological link, i.e., different domain, location, ward/floc and sample.

## Results

### Isolate characteristics

Sixteen of 107 isolates contained both an IncI1 pST12 and a *bla*_CMY-2_ gene (**supplementary table S1**). Based upon the above-mentioned selection criteria, fourteen isolates were included for long-read sequencing analysis, i.e. thirteen *E. coli* and one *Salmonella enterica*, serotype Kentucky (**table 1**.). Nine of the *E. coli* isolates were from one broiler farm, the other isolates were from human origin. The *E. coli* isolates included five different MLSTs and *fim* types. Based on wgMLST analysis four different clusters could identified (**table 1**. and **supplementary table S2**). Additional information regarding antimicrobial resistance phenotype and genotype of the included isolates is provided in **Supplementary table S3**.

**Table 1.** **Descriptive characteristics of** fourteen IncI1 pST12 and *bla*_CMY-2_ containing isolates

| Isolate no. | Species | MultilocusST<sup>a</sup> | wgMLST<br>cluster | Fim | Origin | Sample location | Flock or ward | Sample<br>source | Month and Year of isolation | Accession<br>no. |
| --- | --- | --- | --- | --- | --- | --- | --- | --- | --- | --- |
| EC1 | *E. coli* | ST665 | 1 | fimH30 | Broiler | Farm 1, | Flock 1 | Fecal swab 1 | Nov 2017 | ERS4591617 |
| EC2 | *E. coli* | ST665 | 1 | fimH30 | Broiler | Farm 1 | Flock 1 | Fecal swab 2 | Nov 2017 | ERS4591618 |
| EC3 | *E. coli* | ST665 | 1 | fimH30 | Broiler | Farm 1 | Flock 2 | Fecal swab 3 | Nov 2017 | ERS4591619 |
| EC4 | *E. coli* | ST665 | 1 | fimH30 | Broiler | Farm 1 | Flock 2 | Fecal swab 4 | Nov 2017 | ERS4591620 |
| EC5 | *E. coli* | ST665 | 1 | fimH30 | Broiler | Farm 1 | Flock 3 | Fecal swab 5 | Nov 2017 | ERS4591621 |
| EC6 | *E. coli* | ST665 | 1 | fimH30 | Broiler | Farm 1 | Flock 3 | Fecal swab 6 | Nov 2017 | ERS4591622 |
| EC7 | *E. coli* | ST86 | 2 | fimH289 | Broiler | Farm 1 | Flock 3 | Fecal swab 5 | Nov 2017 | ERS4591623 |
| EC8 | *E. coli* | ST86 | 2 | fimH289 | Broiler | Farm 1 | Flock 3 | Fecal swab 7 | Nov 2017 | ERS4591624 |
| EC9 | *E. coli* | ST6856 |  | fimH71 | Broiler | Farm 1 | Flock 3 | Fecal swab 6 | Nov 2017 | ERS4591625 |
| EC10 | *E. coli* | ST131 | 3 | fimH22 | Human | Hospital 1 | Ward 1 | Blood 1 | Oct 2013 | ERS4591626 |
| EC11 | *E. coli* | ST131 | 3 | fimH22 | Human | Hospital 2 | Ward 1 | Blood 2 | Jul 2014 | ERS4591627 |
| EC12 | *E. coli* | ST973 | 4 | fimH95 | Human | Hospital 3 | Ward 1 | Rectal swab 1 | Dec 2017 | ERS4591628 |
| EC13 | *E. coli* | ST973 | 4 | fimH95 | Human | Hospital 3 | Ward 2 | Rectal swab 2 | Dec 2017 | ERS4591629 |
| SE1 | *Salmonella enteritidis* | - |  | - | Human | Primary care unit | n.a. | Feces | Aug 2018 | ERS4591630 |

a. MLST according to Enterobase (http://enterobase.warwick.ac.uk/)

### Plasmid analysis

In the hybrid assembly of 14 sequences, both the IncI1-pST12 replicon gene and *bla*_CMY-2_ gene were located on a single circular contig ranging in size from 98,410 to 98,999 bp. No additional antimicrobial resistance or virulence genes were detected on any of the extracted plasmids. The number of SNP’s detected between the fourteen plasmids ranged from 0 to 9 SNP’s (**Table 2**). When comparing the plasmids extracted from the selected isolates to a publicly available IncI1-pST12 *bla*_CMY-2_ gene-containing plasmid extracted from the GenBank (accession number: MH472638.1), the number of SNP’s detected ranged from 0 to 7 (**Table 2**). The range of SNP differences overlapped between epidemiologically related and unrelated plasmids (**Table 3**). The median number of SNP differences of plasmids in a different domain or different location, but the same domain, was higher than in the other three pairwise comparison groups.

**Table 2.** Number of SNP’s detected between the 14 extracted plasmids and GenBank reference plasmid MH472638.1.

|  | pEC1 | pEC2 | pEC3 | pEC4 | pEC5 | pEC6 | pEC7 | pEC8 | pEC9 | pEC10 | pEC11 | pEC12 | pEC13 | pSE1 | MH472638.1 |
| --- | --- | --- | --- | --- | --- | --- | --- | --- | --- | --- | --- | --- | --- | --- | --- |
| pEC1 | 0 | 2 | 2 | 2 | 2 | 2 | 3 | 3 | 3 | 3 | 3 | 9 | 8 | 6 | 2 |
| pEC2 | 2 | 0 | 0 | 0 | 0 | 0 | 1 | 1 | 1 | 1 | 1 | 7 | 6 | 4 | 0 |
| pEC3 | 2 | 0 | 0 | 0 | 0 | 0 | 1 | 1 | 1 | 1 | 1 | 7 | 6 | 4 | 0 |
| pEC4 | 2 | 0 | 0 | 0 | 0 | 0 | 1 | 1 | 1 | 1 | 1 | 7 | 6 | 4 | 0 |
| pEC5 | 2 | 0 | 0 | 0 | 0 | 0 | 1 | 1 | 1 | 1 | 1 | 7 | 6 | 4 | 0 |
| pEC6 | 2 | 0 | 0 | 0 | 0 | 0 | 1 | 1 | 1 | 1 | 1 | 7 | 6 | 4 | 0 |
| pEC7 | 3 | 1 | 1 | 1 | 1 | 1 | 0 | 0 | 0 | 2 | 2 | 8 | 7 | 5 | 1 |
| pEC8 | 3 | 1 | 1 | 1 | 1 | 1 | 0 | 0 | 0 | 2 | 2 | 8 | 7 | 5 | 1 |
| pEC9 | 3 | 1 | 1 | 1 | 1 | 1 | 0 | 0 | 0 | 2 | 2 | 8 | 7 | 5 | 1 |
| pEC10 | 3 | 1 | 1 | 1 | 1 | 1 | 2 | 2 | 2 | 0 | 0 | 8 | 7 | 5 | 1 |
| pEC11 | 3 | 1 | 1 | 1 | 1 | 1 | 2 | 2 | 2 | 0 | 0 | 8 | 7 | 5 | 1 |
| pEC12 | 9 | 7 | 7 | 7 | 7 | 7 | 8 | 8 | 8 | 8 | 8 | 0 | 1 | 5 | 7 |
| pEC13 | 8 | 6 | 6 | 6 | 6 | 6 | 7 | 7 | 7 | 7 | 7 | 1 | 0 | 4 | 6 |
| pSE1 | 6 | 4 | 4 | 4 | 4 | 4 | 5 | 5 | 5 | 5 | 5 | 5 | 4 | 0 | 4 |
| MH472638.1 | 2 | 0 | 0 | 0 | 0 | 0 | 1 | 1 | 1 | 1 | 1 | 7 | 6 | 4 | 0 |

**Table 3.** Median and range of SNP differences in pairwise comparisons per EPI link

|  | *n* of pairwise comparisons | SNP differences |  |
| --- | --- | --- | --- |
|  |  | Median | Range |
| Same sample<sup>(i)</sup> | 2 | 1 | 1 |
| Same flock, different sample<sup>(ii)</sup> | 10 | 0.5 | 0-2 |
| Same location, different ward/flock<sup>(iii)</sup> | 25 | 1 | 0-3 |
| Same domain, different location<sup>(iv)</sup> | 9 | 5 | 0-8 |
| Different domain<sup>(v)</sup> | 45 | 4 | 1-9 |

The total number of genes detected in the fourteen plasmids was 113 of which 112 were detected in all plasmids. One gene was present only in one plasmid (pEC11) and encoded for a hypothetical protein. An alignment of coding regions of the fourteen plasmids revealed no rearrangements between the described plsmids (**Supplementary Figure 1)**. However, progressive MAUVE alignment of non-coding regions revealed a small highly-variable region of 519 to 1096bp in all plasmids. This variable region contained insertions, deletions and inversions of four genetic elements. (**Supplementary Figure 2**). No rearrangements were detected in any of the other regions.

## Discussion

The current study included *E. coli* isolates of various sequence types and a *S. enterica* isolate, which were from both human and broiler origin. Plasmid analysis based on short- and long-read sequence data of *bla*_CMY-2_ containing IncI1-pST12 plasmids from the included isolates revealed a low number of SNP differences and a high number of shared genes between the various plasmids extracted. Despite the tendency of median SNP increase from epidemiologically related to unrelated plasmids, the range in number of SNPs detected overlapped between every classified epidemiological link in the current study. Furthermore, only one gene was variably present between the different plasmids and no rearrangements were observed apart from a small, highly variable region. Probably this area is the formerly described highly variable shufflon region at the C-terminal end of the PilV protein [34, 35].

A high degree of similarity between IncI1-pST12 plasmids was previously reported [12–14]. However, these studies either contained only plasmids extracted from one *E. coli* sequence type (ST131) [12], or included plasmids were primarily of poultry origin [14]. Moreover, these studies predominately used *in silico* reference-based plasmid reconstructions of short-read sequence data rather than performing a hybrid assembly of both short- and long-read sequence data. A recent study by Valcek *et al*. on IncI1-pST3 and -pST7 plasmids showed that using combined long-read and short-read sequencing data improves the accuracy of a full plasmid analysis, e.g. of rearrangements [15]. All of studies used either gene presence/absence based or SNP based analysis, but not both, possibly missing subtle differences between various plasmids.

Several studies have described outbreaks with *bla*_CMY-2_ harbouring Enterobacteriaceae [36–39]. Since the *bla*_CMY-2_ is predominantly located on plasmids, horizontal transfer of the plasmid in an outbreak can go undetected if only typing of the bacterial chromosome is performed. Distinguishing epidemiologically related and unrelated plasmids is essential to confirm plasmid transmission in an outbreak. Therefore, statements on horizontal transfer of these plasmids based on genetic identity should be made with caution.

The current study is the first to explore *bla*_CMY-2_ containing IncI1-pST12 plasmids from related and unrelated isolates, using combined short- and long-read sequencing data. Moreover, this study includes isolates from different species, sequence types and domains, both from human and broiler origin. Two different comparison techniques, either gene presence/absence and SNP differences, were used. Furthermore, combining of long-read and short-read sequence data provided full plasmid analysis, including the presence of rearrangements.

A limitation of the current study is that the small sample size precludes the use of statistical test and caution must be applied, as the findings should be confirmed in a study with a larger sample size. Preferably, such a study should include isolates of different species, sequence types, and origin of isolation containing IncI1-pST12 plasmids. Furthermore, the current study only included plasmids of broilers isolated in one farm, therefore other plasmids of veterinary origin should be added in future studies to confirm our findings.

In conclusion, IncI1-pST12 plasmids of epidemiologically related and unrelated Enterobacteriaceae of both human and broiler origin in the current explorative study show a high degree of sequence similarity in terms of SNP differences and the number of shared genes.

## Acknowledgements

We are grateful to the collaborators from the in the participating laboratories, hospitals and livestock farms for their contribution to the collection of the microbiological and epidemiological data.

## i-4-1-Health Study Group

Lieke van Alphen (Maastricht University Medical Center+, Maastricht, the Netherlands), Nicole van den Braak (Avans University of Applied Sciences, Breda, the Netherlands), Caroline Broucke (Agency for Care and Health, Brussels, Belgium), Anton Buiting (Elisabeth-TweeSteden Hospital, Tilburg, the Netherlands), Liselotte Coorevits (Ghent University Hospital, Ghent, Belgium), Sara Dequeker (Agency for Care and Health, Brussels, Belgium and Sciensano, Brussels, Belgium), Jeroen Dewulf (Ghent University, Ghent, Belgium), Wouter Dhaeze (Agency for Care and Health, Brussels, Belgium), Bram Diederen (ZorgSaam Hospital, Terneuzen, the Netherlands), Helen Ewalts (Regional Public Health Service Hart voor Brabant, Tilburg, the Netherlands), Herman Goossens (University of Antwerp, Antwerpen, Belgium and Antwerp University Hospital, Antwerp, Belgium), Inge Gyssens (Hasselt University, Hasselt, Belgium), Casper den Heijer (Regional Public Health Service ZuidLimburg, Heerlen, the Netherlands), Christian Hoebe (Maastricht University Medical Center+, Maastricht, the Netherlands and Regional Public Health Service Zuid-Limburg, Heerlen, Casper Jamin (Maastricht University Medical Center+, Maastricht, the Netherlands), Patricia Jansingh (Regional Public Health Service Limburg Noord, Venlo, the Netherlands), Jan Kluytmans (Amphia Hospital, Breda, the Netherlands and University Medical Center Utrecht, Utrecht University, Utrecht, the Netherlands), Marjolein Kluytmans-van den Bergh (Amphia Hospital, Breda, the Netherlands and University Medical Center Utrecht, Utrecht University, Utrecht, the Netherlands), Stefanie van Koeveringe (Antwerp University Hospital, Antwerp, Belgium), Sien De Koster (University of Antwerp, Antwerp, Belgium), Christine Lammens (University of Antwerp, Antwerp, Belgium), Isabel Leroux-Roels (Ghent University Hospital, Ghent, Belgium), Hanna Masson (Agency for Care and Health, Brussel, Belgium), Ellen Nieuwkoop (Elisabeth-TweeSteden Hospital, Tilburg, the Netherlands), Anita van Oosten (Admiraal De Ruyter Hospital, Goes, the Netherlands), Natascha Perales Selva (Antwerp University Hospital, Antwerp, Belgium), Merel Postma (Ghent University, Ghent, Belgium), Stijn Raven (Regional Public Health Service West-Brabant, Breda, the Netherlands), Veroniek Saegeman (University Hospitals Leuven, Leuven, Belgium), Paul Savelkoul (Maastricht University Medical Center+, Maastricht, the Netherlands), Annette Schuermans (University Hospitals Leuven, Leuven, Belgium), Nathalie Sleeckx (Experimental Poultry Centre, Geel, Belgium), Arjan Stegeman (Utrecht University, Utrecht, the Netherlands), Tijs Tobias (Utrecht University, Utrecht, the Netherlands), Paulien Tolsma (Regional Public Health Service Brabant Zuid-Oost, Eindhoven, the Netherlands), Jacobien Veenemans (Admiraal De Ruyter Hospital, Goes, the Netherlands), Dewi van der Vegt (PAMM Laboratory for Pathology and Medical Microbiology, Veldhoven, the Netherlands), Francisca Velkers (Utrecht University, Utrecht, the Netherlands), Martine Verelst (University Hospitals Leuven, Leuven, Belgium), Carlo Verhulst (Amphia Hospital, Breda, the Netherlands), Pascal De Waegemaeker (Ghent University Hospital, Ghent, Belgium), Veronica Weterings (Amphia Hospital, Breda, the Netherlands), Clementine Wijkmans (Regional Public Health Service Hart voor Brabant, Tilburg, the Netherlands), Patricia Willemse-Smits (Elkerliek Hospital, Helmond, the Netherlands), Ina Willemsen (Amphia Hospital, Breda, the Netherlands).

## Transparency declarations

None to declare.

## Supplementary data

- Supplementary table 1
- Supplementary table 2
- Supplementary table 3
- Supplementary Figure 1
- Supplementary Figure 2

## References

1. Davies J. Origins and evolution of antibiotic resistance. Microbiologia 1996;12:9–16.

2. Premanandh J, Samara BS, Mazen AN. Race Against Antimicrobial Resistance Requires Coordinated Action – An Overview. Front Microbiol 2016;6:1–6.

3. Carlet J. The gut is the epicentre of antibiotic resistance. Antimicrob Resist Infect Control 2012;1:1–7.

4. Carattoli A. Animal reservoirs for extended spectrum β-lactamase producers. Clin Microbiol Infect 2008;14:117–123.

5. Ewers C, Bethe A, Semmler T, Guenther S, Wieler LH. Extended-spectrum β-lactamase-producing and AmpC-producing Escherichia coli from livestock and companion animals, and their putative impact on public health: A global perspective. Clin Microbiol Infect 2012;18:646–655.

6. Dierikx C, van der Goot J, Fabri T, van Essen-Zandbergen A, Smith H, et al. Extended-spectrum-β-lactamase- and AmpC-β-lactamase-producing Escherichia coli in Dutch broilers and broiler farmers. J Antimicrob Chemother 2013;68:60–67.

7. Berg ES, Wester AL, Ahrenfeldt J, Mo SS, Slettemeås JS, et al. Norwegian patients and retail chicken meat share cephalosporin-resistant Escherichia coli and IncK/blaCMY-2 resistance plasmids. Clin Microbiol Infect 2017;23:407.e9-407.e15.

8. Jacoby GA. AmpC B-Lactamases. Clin Microbiol Rev 2009;22:161–182.

9. Accogli M, Fortini D, Giufrè M, Graziani C, Dolejska M, et al. IncI1 plasmids associated with the spread of CMY-2, CTX-M-1 and SHV-12 in Escherichia coli of animal and human origin. Clin Microbiol Infect 2013;19:E238–40.

10. Hansen KH, Bortolaia V, Nielsen CA, Nielsen JB, Schønning K, et al. Host-specific patterns of genetic diversity among IncI1-I? and IncK plasmids encoding CMY-2 β-lactamase in escherichia coli isolates from humans, poultry meat, poultry, and dogs in Denmark. Appl Environ Microbiol 2016;82:4705–4714.

11. Carattoli A, Villa L, Fortini D, García-Fernández A. Contemporary IncI1 plasmids involved in the transmission and spread of antimicrobial resistance in Enterobacteriaceae. Plasmid. Epub ahead of print December 2018. DOI: 10.1016/j.plasmid.2018.12.001.

12. Roer L, Overballe-Petersen S, Hansen F, Johannesen TB, Stegger M, et al. ST131 fimH22 Escherichia coli isolate with a blaCMY-2/IncI1/ST12 plasmid obtained from a patient with bloodstream infection: Highly similar to E. coli isolates of broiler origin. J Antimicrob Chemother 2019;74:557–560.

13. Pietsch M, Irrgang A, Roschanski N, Brenner Michael G, Hamprecht A, et al. Whole genome analyses of CMY-2-producing Escherichia coli isolates from humans, animals and food in Germany. BMC Genomics;19. Epub ahead of print 2018. DOI: 10.1186/s12864-018-4976-3.

14. Castellanos LR, van der Graaf-Van Bloois L, Donado-Godoy P, Mevius DJ, Wagenaar JA, et al. Phylogenomic investigation of IncI1-I plasmids harboring blaCMY-2 and blaSHV-12 in salmonella enterica and Escherichia coli in multiple countries. Antimicrob Agents Chemother;63. Epub ahead of print 2019. DOI: 10.1128/AAC.02546-18.

15. Valcek A, Roer L, Overballe-Petersen S, Hansen F, Bortolaia V, et al. IncI1 ST3 and IncI1 ST7 plasmids from CTX-M-1-producing Escherichia coli obtained from patients with bloodstream infections are closely related to plasmids from E. coli of animal origin. J Antimicrob Chemother 2019;74:2171–2175.

16. Stohr JJJM, Bergh MFQK den, Wedema R, Friedrich AW, Kluytmans JAJW, et al. Detection of extended-spectrum beta-lactamase (ESBL) genes and plasmid replicons in Enterobacteriaceae using PlasmidSPAdes assembly of short-read sequence data. bioRxiv 2019;863316.

17. Bergh MK Den, Lammens C, Selva NP, Buiting A, Leroux-roels I, et al. Microbiological methods to detect intestinal carriage of highly-resistant microorganisms (HRMO) in humans and livestock in the i-4-1-Health Dutch-Belgian cross-border project. Preprints.org 2019;1–16.

18. Drijver E den, Verweij JJ, Verhulst C, Soer J, Veldman K, et al. Detection of AmpC β-lactamases in Escherichia coli using different screening Evert den Drijver. BioRxiv. Epub ahead of print 2019. DOI: http://dx.doi.org/10.1101/787085.

19. Coolen JPM, Den Drijver EPM, Kluytmans JAJW, Verweij JJ, Lamberts BA, et al. Development of an algorithm to discriminate between plasmid- and chromosomal-mediated AmpC β-lactamase production in Escherichia coli by elaborate phenotypic and genotypic characterization. J Antimicrob Chemother 2019;74:3481–3488.

20. Bankevich A, Nurk S, Antipov D, Gurevich AA, Dvorkin M, et al. SPAdes: A New Genome Assembly Algorithm and Its Applications to Single-Cell Sequencing. J Comput Biol 2012;19:455–477.

21. Den Drijver E, Verweij JJ, Verhulst C, Oome S, Soer J, et al. Decline in AmpC β-lactamase-producing escherichia coli in a Dutch teaching hospital (2013-2016). PLoS One;13. Epub ahead of print 2018. DOI: 10.1371/journal.pone.0204864.

22. Kluytmans-Van Den Bergh MFQ, Rossen JWA, Bruijning-Verhagen PCJ, Bonten MJM, Friedrich AW, et al. Whole-genome multilocus sequence typing of extended-spectrum-beta-lactamase-producing enterobacteriaceae. J Clin Microbiol 2016;54:2919–2927.

23. Zankari E, Hasman H, Cosentino S, Vestergaard M, Rasmussen S, et al. Identification of acquired antimicrobial resistance genes. J Antimicrob Chemother 2012;67:2640–2644.

24. Carattoli A, Zankari E, Garciá-Fernández A, Larsen MV, Lund O, et al. In Silico detection and typing of plasmids using plasmidfinder and plasmid multilocus sequence typing. Antimicrob Agents Chemother 2014;58:3895–3903.

25. Larsen M V., Cosentino S, Rasmussen S, Friis C, Hasman H, et al. Multilocus sequence typing of total-genome-sequenced bacteria. J Clin Microbiol 2012;50:1355–1361.

26. Camacho C, Coulouris G, Avagyan V, Ma N, Papadopoulos J, et al. BLAST+: Architecture and applications. BMC Bioinformatics;10. Epub ahead of print 2009. DOI: 10.1186/1471-2105-10-421.

27. Wick RR, Judd LM, Gorrie CL, Holt KE. Unicycler: Resolving bacterial genome assemblies from short and long sequencing reads. PLoS Comput Biol 2017;13:e1005595.

28. Joensen KG, Scheutz F, Lund O, Hasman H, Kaas RS, et al. Real-time whole-genome sequencing for routine typing, surveillance, and outbreak detection of verotoxigenic Escherichia coli. J Clin Microbiol 2014;52:1501–1510.

29. Wick RR, Schultz MB, Zobel J, Holt KE. Bandage: Interactive visualization of de novo genome assemblies. Bioinformatics 2015;31:3350–3352.

30. Seemann T. Prokka: Rapid prokaryotic genome annotation. Bioinformatics 2014;30:2068–2069.

31. Page AJ, Cummins CA, Hunt M, Wong VK, Reuter S, et al. Roary: Rapid large-scale prokaryote pan genome analysis. Bioinformatics 2015;31:3691–3693.

32. Petkau A, Stuart-Edwards M, Stothard P, van Domselaar G. Interactive microbial genome visualization with GView. Bioinformatics 2010;26:3125–3126.

33. Darling AE, Mau B, Perna NT. Progressivemauve: Multiple genome alignment with gene gain, loss and rearrangement. PLoS One;5. Epub ahead of print 2010. DOI: 10.1371/journal.pone.0011147.

34. Brouwer MSM, Tagg KA, Mevius DJ, Iredell JR, Bossers A, et al. IncI shufflons: Assembly issues in the next-generation sequencing era. Plasmid 2015;80:111–117.

35. Brouwer MSM, Jurburg SD, Harders F, Kant A, Mevius DJ, et al. The shufflon of IncI1 plasmids is rearranged constantly during different growth conditions. Plasmid 2019;102:51–55.

36. Wendorf KA, Kay M, Baliga C, Weissman SJ, Gluck M, et al. Endoscopic Retrograde Cholangiopancreatography-Associated AmpC Escherichia coli Outbreak. Infect Control Hosp Epidemiol 2015;36:634–642.

37. Huang IF, Chiu CH, Wang MH, Wu CY, Hsieh KS, et al. Outbreak of dysentery associated with ceftriaxone-resistant Shigella sonnei: First report of plasmid-mediated CMY-2-type AmpC β-lactamase resistance in S. sonnei. J Clin Microbiol 2005;43:2608–2612.

38. Kameyama M, Yabata J, Nomura Y, Tominaga K. Detection of CMY-2 AmpC β-lactamase-producing enterohemorrhagic Escherichia coli O157: H7 from outbreak strains in a nursery school in Japan. J Infect Chemother 2015;21:544–546.

39. Matsumura Y, Tanaka M, Yamamoto M, Nagao M, Machida K, et al. High prevalence of carbapenem resistance among plasmid-mediated AmpC β-lactamase-producing Klebsiella pneumoniae during outbreaks in liver transplantation units. Int J Antimicrob Agents 2015;45:33–40.

