## Supplementary Table 1 for "Limited genetic diversity of *bla*_CMY-2_-containing IncI1-pST12 plasmids from Enterobacteriaceae of human and broiler chicken origin in the Netherlands"

Supplementary table S1. Number of sequenced isolates in various collections, isolates containing a IncI1-pST12 replicon and blaCMY-2 gene, and included isolates.

| Collection | N of isolates | | N of isolates containing IncI1-pST12 replicon and *bla* $_{CMY-2}$ gene | | N of isolates included in study | |
|---|---|---|---|---|---|---|
| | Human | Broiler | Human | Broiler | Human | Broiler |
| I-4-1 Health | 19 | 22 | 0 | 11 | 0 | 9 |
| Amphia prevalence screening | 14 | 0 | 2 | 0 | 2 | 0 |
| Blood cultures | 51 | 0 | 2 | 0 | 2 | 0 |
| Fecal culture | 1 | 0 | 1 | 0 | 1 | 0 |
