## Supplementary table 2 for "Limited genetic diversity of *bla*_CMY-2_-containing IncI1-pST12 plasmids from Enterobacteriaceae of human and broiler chicken origin in the Netherlands"

Supplementary table S2. Distance matrix containing number of allele diffferences between the included *E. coli* isolates based on wgMLST

| | EC1 | EC2 | EC3 | EC4 | EC5 | EC6 | EC7 | EC8 | EC9 | EC10 | EC11 | EC12 | EC13 |
|---|---|---|---|---|---|---|---|---|---|---|---|---|---|
| EC1 | 0 | 0 | 0 | 0 | 0 | 0 | 2051 | 2051 | 1369 | 2318 | 2324 | 2288 | 2286 |
| EC2 | 0 | 0 | 0 | 0 | 0 | 0 | 2053 | 2053 | 1370 | 2320 | 2326 | 2290 | 2288 |
| EC3 | 0 | 0 | 0 | 0 | 0 | 0 | 2050 | 2050 | 1367 | 2317 | 2323 | 2287 | 2285 |
| EC4 | 0 | 0 | 0 | 0 | 0 | 0 | 2051 | 2051 | 1368 | 2318 | 2324 | 2288 | 2286 |
| EC5 | 0 | 0 | 0 | 0 | 0 | 0 | 2053 | 2053 | 1370 | 2320 | 2326 | 2290 | 2288 |
| EC6 | 0 | 0 | 0 | 0 | 0 | 0 | 2053 | 2053 | 1370 | 2320 | 2326 | 2290 | 2288 |
| EC7 | 2051 | 2053 | 2050 | 2051 | 2053 | 2053 | 0 | 1 | 2021 | 2338 | 2344 | 2304 | 2302 |
| EC8 | 2051 | 2053 | 2050 | 2051 | 2053 | 2053 | 1 | 0 | 2021 | 2338 | 2344 | 2304 | 2302 |
| EC9 | 1369 | 1370 | 1367 | 1368 | 1370 | 1370 | 2021 | 2021 | 0 | 2273 | 2279 | 2244 | 2242 |
| EC10 | 2318 | 2320 | 2317 | 2318 | 2320 | 2320 | 2338 | 2338 | 2273 | 0 | 1 | 2363 | 2361 |
| EC11 | 2324 | 2326 | 2323 | 2324 | 2326 | 2326 | 2344 | 2344 | 2279 | 1 | 0 | 2369 | 2367 |
| EC12 | 2288 | 2290 | 2287 | 2288 | 2290 | 2290 | 2304 | 2304 | 2244 | 2363 | 2369 | 0 | 7 |
| EC13 | 2286 | 2288 | 2285 | 2286 | 2288 | 2288 | 2302 | 2302 | 2242 | 2361 | 2367 | 7 | 0 |
