## Supplementary table 3 for "Limited genetic diversity of *bla*_CMY-2_-containing IncI1-pST12 plasmids from Enterobacteriaceae of human and broiler chicken origin in the Netherlands"

**Supplementary table 3.** Overview of resistance genes and phenotype of  the fourteen IncI1 pST12 and bla$_{CMY-2}$ containing isolates

| Isolate no. | Species | MultilocusST[a] | Resistance genes | Piperacillin-Tazobactam MIC | Cefotaxime/Ceftriaxone MIC | Ceftazidime MIC | Cefoxitin MIC | Accession no.[b] |
|---|---|---|---|---|---|---|---|---|
| **EC1** | *E. coli* | ST665 | *bla*$_{CMY-2}$, *bla*$_{TEM-1b,}$ *aada1, strA, strB, sul1, sul2, dfra1* | 8 mg/L | 4 mg/L | 32 mg/L | ≥64 mg/L | ERS4591617 |
| **EC2** | *E. coli* | ST665 | *bla*$_{CMY-2}$, *bla*$_{TEM-1b,}$ *aada1, strA, strB, sul1, sul2, dfra1* | 8 mg/L | 8 mg/L | 32 mg/L | ≥64 mg/L | ERS4591618 |
| **EC3** | *E. coli* | ST665 | *bla*$_{CMY-2}$, *bla*$_{TEM-1b,}$ *aada1, strA, strB, sul1, sul2, dfra1* | 8 mg/L | 4 mg/L | 32 mg/L | ≥64 mg/L | ERS4591619 |
| **EC4** | *E. coli* | ST665 | *bla*$_{CMY-2}$*aada1, strA, strB, sul1, sul2, dfra1* | 8 mg/L | 8 mg/L | 32 mg/L | ≥64 mg/L | ERS4591620 |
| **EC5** | *E. coli* | ST665 | *bla*$_{CMY-2}$, *bla*$_{TEM-1b,}$ *aada1, strA, strB, sul1, sul2, dfra1* | 8 mg/L | 4 mg/L | 32 mg/L | ≥64 mg/L | ERS4591621 |
| **EC6** | *E. coli* | ST665 | *bla*$_{CMY-2}$, *bla*$_{TEM-1b,}$ *aada1, strA, strB, sul1, sul2, dfra1* | 8 mg/L | 4 mg/L | 32 mg/L | ≥64 mg/L | ERS4591622 |
| **EC7** | *E. coli* | ST86 | *bla*$_{CMY-2}$, *bla*$_{TEM-1b, ,}$ *strA, strB, sul2, dfra1* | ≤4 mg/L | 4 mg/L | 8 mg/L | 32 mg/L | ERS4591623 |
| **EC8** | *E. coli* | ST86 | *bla*$_{CMY-2}$, *bla*$_{TEM-1b,}$ *strA, strB, sul2, dfra1* | ≤4 mg/L | 8 mg/L | 8 mg/L | 32 mg/L | ERS4591624 |
| **EC9** | *E. coli* | ST6856 | *bla*$_{CMY-2}$, *bla*$_{TEM-1b,}$ | 8 mg/L | 8 mg/L | 32 mg/L | ≥64 mg/L | ERS4591625 |
| **EC10** | *E. coli* | ST131 | *bla*$_{CMY-2}$, *bla*$_{TEM-1b,}$ *QnrS1, aac(6')-Ib-cr, aadA16-like*[c], *ARR-3, sul1, dfrA27, tet(A)-like*[c] | ≥128 mg/L | ≥64 mg/L | ≥64 mg/L | ≥64 mg/L | ERS4591626 |
| **EC11** | *E. coli* | ST131 | *bla*$_{CMY-2}$, *bla*$_{TEM-1b,}$ *QnrS1* | 8 mg/L | 8 mg/L | 16 mg/L | ≥64 mg/L | ERS4591627 |
| **EC12** | *E. coli* | ST973 | *bla*$_{CMY-2}$ | 8 mg/L | >64 mg/L | 32 mg/L | ≥64 mg/L | ERS4591628 |
| **EC13** | *E. coli* | ST973 | *bla*$_{CMY-2}$ | 8 mg/L | 8 mg/L | 32 mg/L | ≥64 mg/L | ERS4591629 |
| **SE1** | *Salmonella enteritidis* | n.a | *bla*$_{CMY-2}$, *bla*$_{TEM-1b,}$ *aac(6')-Iaa-like, rmtB, catA2-like*[c], *tet(A)-like*[c] | 8 mg/L | >4 mg/L | >16 mg/L | >16 mg/L | ERS4591630 |

a. MLST according to Enterobase (http://enterobase.warwick.ac.uk/)

b. Accession numbers provided by the European Nucleotide Archive (ENA, https://www.ebi.ac.uk/ena)

c. Based on ResFinder version 3.1 Identity score <100% (cge.cbs.dtu.dk/services/ResFinder)
