## Supplementary figures and images for "Limited genetic diversity of *bla*_CMY-2_-containing IncI1-pST12 plasmids from Enterobacteriaceae of human and broiler chicken origin in the Netherlands"

### Supplementary Figure 1

#### IncI1 pST12 CMY-II plasmids

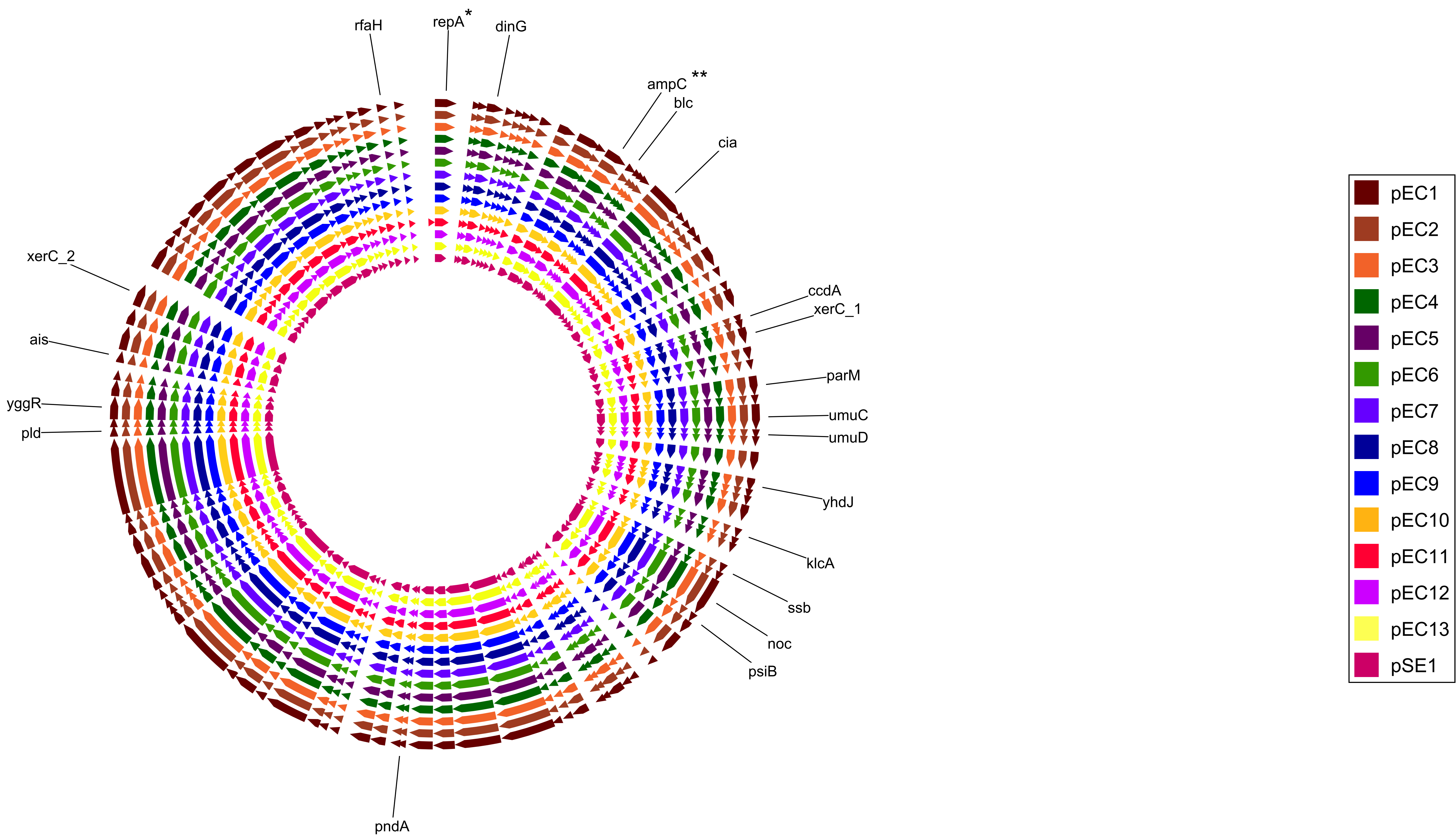


*** repA = IncI1-pST12 replicon  ** ampC = blaCMY-2 gene**

### Supplementary Figure 2

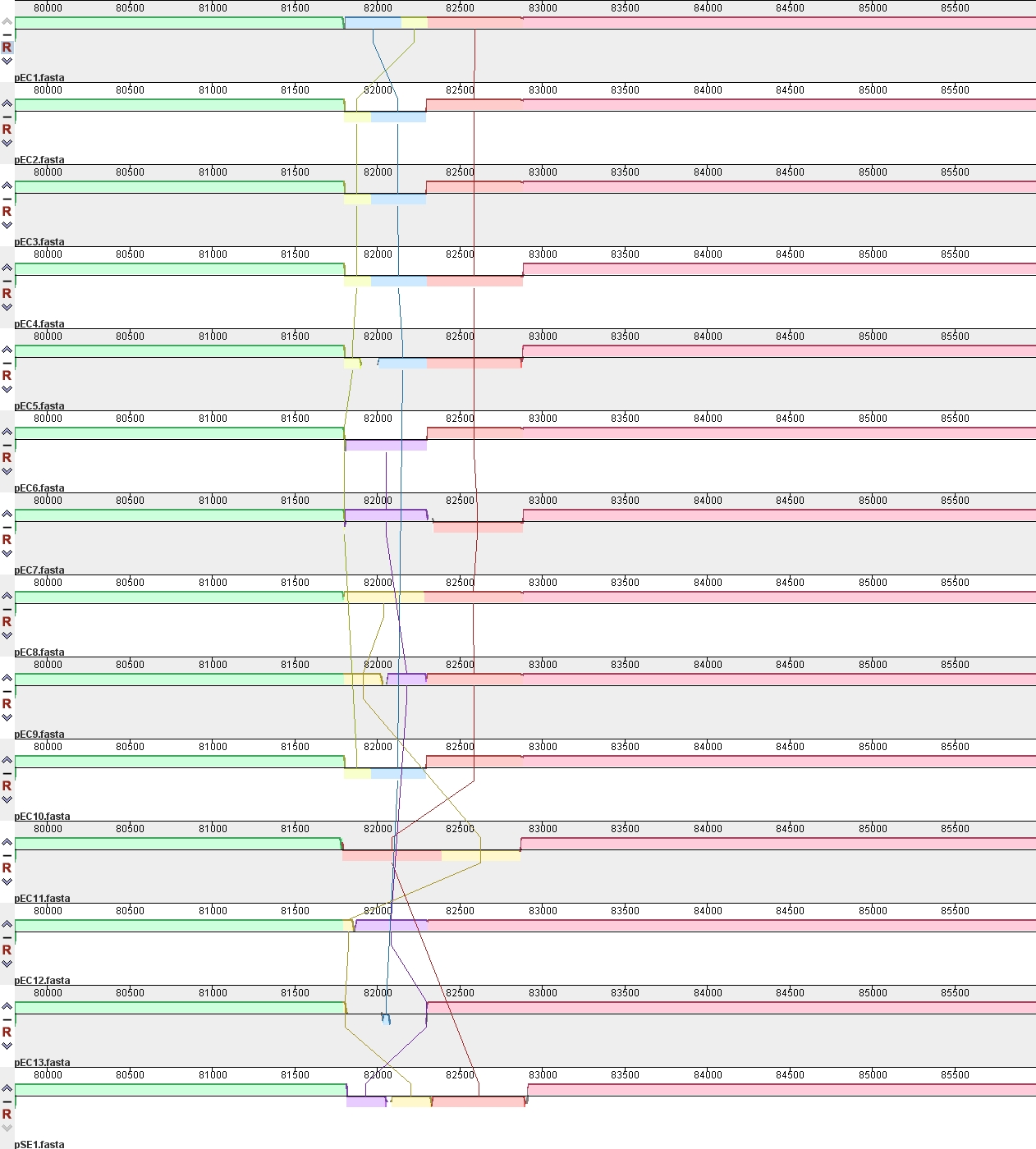
